# Evolutionary trajectories of IDH-mutant astrocytoma identify molecular grading markers related to cell cycling

**DOI:** 10.1101/2024.03.05.583306

**Authors:** Wies R. Vallentgoed, Youri Hoogstrate, Karin A. van Garderen, Levi van Hijfte, Erik van Dijk, Mathilde C. M. Kouwenhoven, Johanna M. Niers, Kaspar Draaisma, Ivonne Martin, Wendy W. J. de Leng, C. Mircea S. Tesileanu, Iris de Heer, Maud Diepeveen, Anna Lavrova, Paul P. Eijk, Marcel Bühler, Wolfgang Wick, Paul M. Clement, Marc Sanson, Enrico Franceschi, Thierry Gorlia, Vassilis Golfinopoulos, Michael Weller, Tobias Weiss, Pierre A. Robe, Johan M. Kros, Marion Smits, Mark van de Wiel, Bauke Ylstra, Roel G. W. Verhaak, Martin J. van den Bent, Bart A. Westerman, Pieter Wesseling, Pim J. French

**Author notes:** Authors contributed equally to this work (Youri Hoogstrate, Karin A. van Garderen, Levi van Hijfte, Erik van Dijk, Bart A. Westerman, Pieter Wesseling, Pim J. French).

## Abstract

To study the evolutionary processes that drive malignant progression of IDH-mutant astrocytomas, we performed multi-omics on a large cohort of matched initial and recurrent tumor samples. The overlay of genetic, epigenetic, transcriptomic and proteomic data, combined with single-cell analysis, have identified overlapping features associated with malignant progression. These features are derived from three molecular mechanisms and provide a rationale of the underlying biology of tumor malignancy: cell-cycling, tumor cell (de-)differentiation and remodeling of the extracellular matrix. Specifically, DNA-methylation levels decreased over time, predominantly in tumors with malignant transformation and co-occurred with poor prognostic genetic events. DNA-methylation was lifted from specific loci associated with DNA replication and was associated with an increased RNA and protein expression of cell cycling associated genes. All results were validated on samples of newly diagnosed IDH-mutant astrocytoma patients included the CATNON randomized phase 3 clinical trial. Importantly, malignant progression was hardly affected by radio- or chemotherapy, indicating that treatment does not affect the course of disease. Our results culminate in a DNA-methylation based signature for objective tumor grading.

## 2 Introduction

Adult-type diffuse gliomas are the most common type of malignant brain tumors and are histologically characterized by their widespread infiltrative properties, without a curative treatment available to date[1]. The classification of central nervous system (CNS) tumors by the World Health Organization (WHO) of 2021 (CNS5), separates these gliomas into IDH-wildtype glioblastomas, and IDH-mutant astrocytomas and oligodendrogliomas[2].

Though molecular characterization has greatly improved our understanding of diffuse gliomas, the processes that drive malignant progression remain poorly understood[3–5]. Even with optimal treatment, consisting of maximal safe resection followed by radio- and/or chemotherapy, these tumors almost always relapse[6]. This tendency towards relapse and malignant transformation makes them ultimately fatal, yet with highly variable survival periods. The Glioma Longitudinal Analysis (GLASS) consortium was initiated to investigate glioma evolution and expose therapeutic vulnerabilities in each of the three major diffuse glioma subtypes[7–9].

Particularly in IDH-mutant astrocytomas, the onset of progression and/or malignant transformation is highly unpredictable. Some of these tumors will relapse particularly early or transform rapidly into a high-grade (CNS WHO grade 4) tumor, despite prior lower (CNS WHO grade 2/3) histological grading. We have therefore initiated the GLASS-NL consortium to comprehensively study the molecular evolution in IDH-mutant, 1p/19q non-codeleted astrocytomas using multi-omics on a large cohort of paired initial and recurrent tumor samples, with extensive clinical annotation.

## 3 Results

### 3.1 Patient and sample characteristics

We identified 111 patients, operated on between 1989 and 2019, with a primary diagnosis of IDH-mutant astrocytoma grade 2-4, based on the WHO CNS5. The final GLASS-NL dataset consists of matched initial and recurrent multi-omics data of 105 patients (**Fig. 1a**). Patient characteristics are summarized in **Table 1**. Tumor material of 3rd and 4th surgical resections were available for 23 and three patients respectively. The median age at diagnosis of this cohort was 32.0 (18.0-70.0) years, which is younger than the average age at diagnosis of grade 2 and 3 IDH-mutant astrocytomas (45 and 52 years respectively)[10]. Treatment regimen were not uniform in this cohort, and additional treatments (i.e. radio- and/or chemotherapy) were not always given adjuvant to a surgical intervention, but also in response to tumor progression. Prior to surgical resection of the initial tumor, six patients underwent a biopsy of the tumor, and two received additional treatment. Over the course of their disease progression, 95 patients received any type of treatment besides surgical resection of the tumor.

**Fig1:**
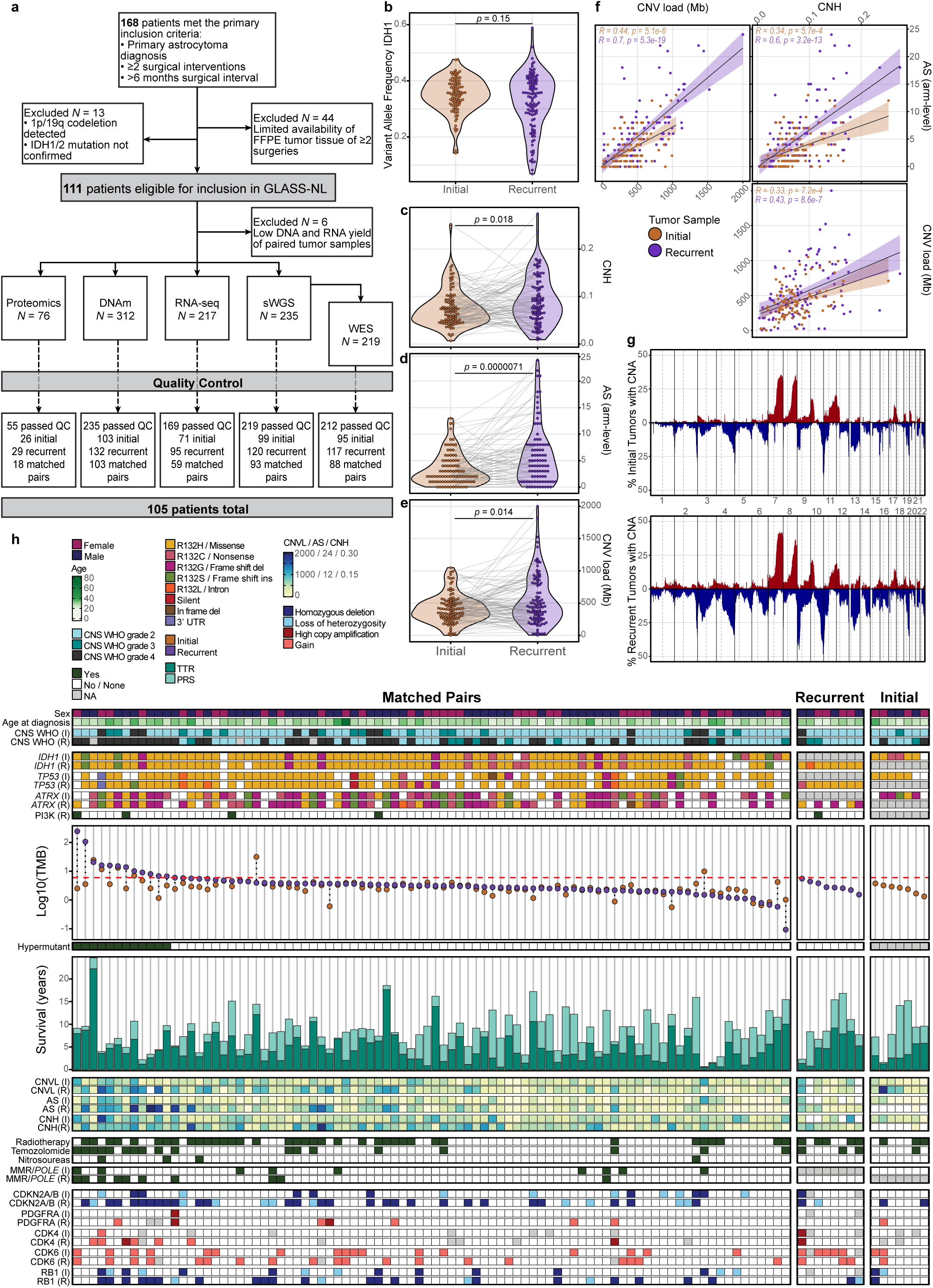
Longitudinal genomic alterations in IDH-mutant astrocytomas. **a**, Flowchart on patient inclusion and data collection of the GLASS-NL cohort. **b**, Difference in tumor purity between initial and recurrent tumor samples based on the VAF of the *IDH1* mutation (see also **Extended Data Fig. 1a**), p-values are calculated using a Wilcoxon rank sum test. **c-e**, Temporal changes in CNH, determined from a single copy-number profile (ranging from 0.004 to 0.274) (**c**), AS, as the sum of altered arms (ranging from 0 to 24) (**d**), and total CNV-load, as the total length of segment-level gains or deletions (ranging from 1Mb-2000Mb) (**e**). Lines connect matched pairs, p-values are calculated using a paired Wilcoxon signed-rank test. **f**, Spearman correlation between AS, CNV-load and CNH in initial and recurrent tumor samples separately (with 95% CI). **g**, Copy-number alteration frequencies (±0.2 Log2(read counts)) in initial (top) and recurrent (bottom) tumor samples, based on sWGS data. **h**, Genomic landscape of the GLASS-NL cohort. Each column represents a patient, the horizontal panels indicate patients with available WES data of (from left to right) a matched tumor pair (n = 88), of only the recurrent tumor (n = 8), and of only the initial tumor (n = 7). Columns are ordered based on the TMB (muts/Mb) of the recurrent tumor, or that of the single tumor sample when there was no matched pair. The vertical panels show (from top to bottom) patient characteristics, mutation type in most commonly altered genes in astrocytoma (*IDH1, TP53, ATRX* and *PIK3CA/PIK3R1* ), TMB of matched tumor pairs with hypermutation cutoff (*≥*6 muts/Mb), patient survival (in TTR and PRS), genomic instability measures (CNH, AS and CNV-load), treatment regimens between initial and recurrent surgery, mutations in MMR genes and/or POLE, CNAs of commonly affected genes in astrocytoma (*CDKN2A/B, PDGFRA, CDK4, CDK6* and *RB1* ). I = initial, R = recurrent, TMB = tumor mutational burden, TTR = time-to-recurrence, PRS = post-recurrent survival, CNVL = total CNV-load, AS = aneuploidy score, CNH = copy-number heterogeneity, MMR = mismatch repair, CNA = copy-number alteration.

**Table 1:**
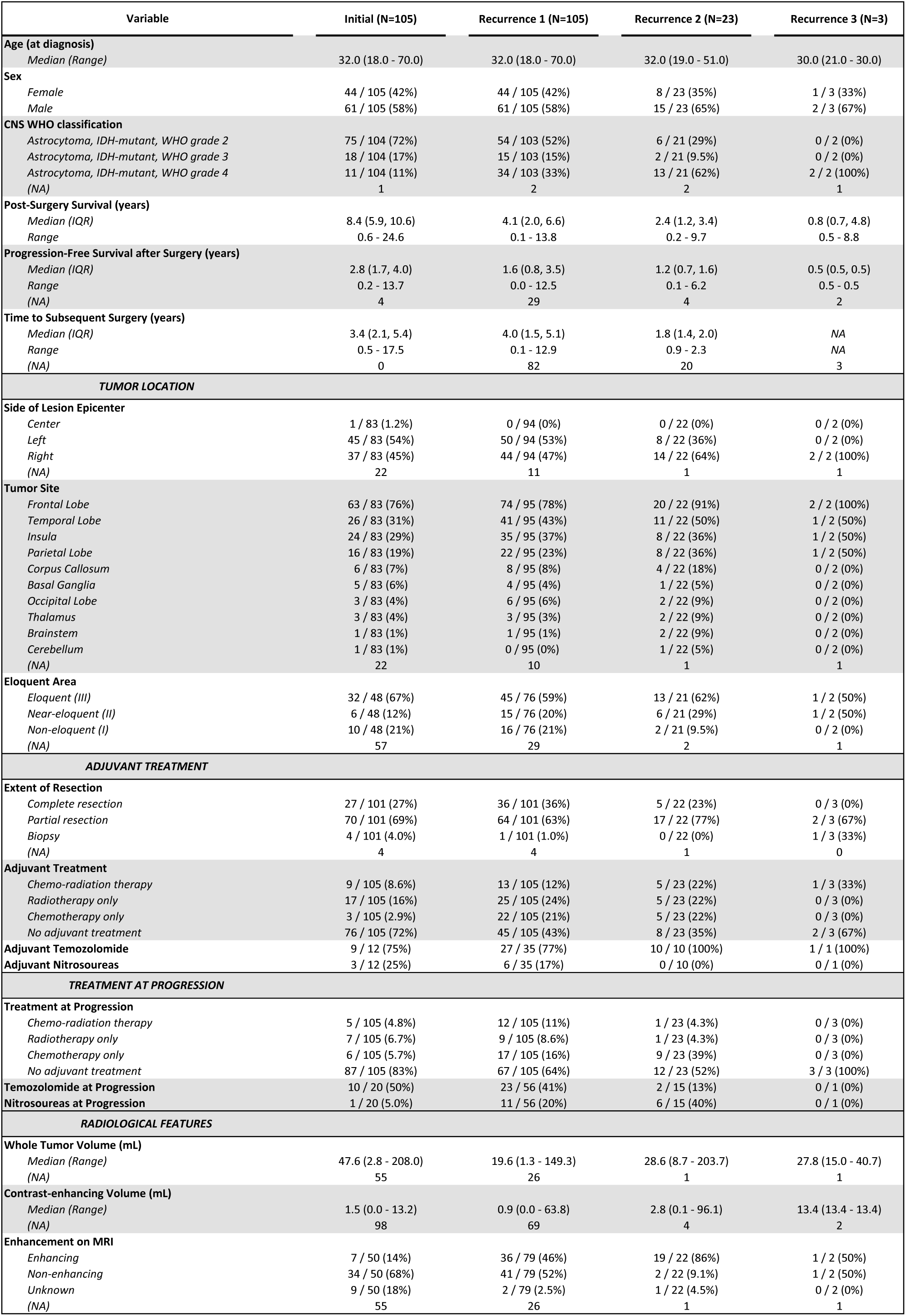
Patient demographic and clinical characteristics of the GLASS-NL cohort. Baseline table of patient, tumor and treatment characteristics for each surgical intervention (initial, n = 105; 1st recurrence, n = 105; 2nd recurrence, n = 23; 3rd recurrence, n = 3) included in the GLASS-NL cohort.

Prior to molecular analyses, we fully curated the available clinical data and analyzed them for their contribution to survival per surgical resection. Age at diagnosis and chemotherapy after initial surgery were independent prognostic factors associated with overall survival (OS), and preoperative contrast-enhancement was associated with post-recurrent survival (PRS) (**Table 2**).

**Table 2:**
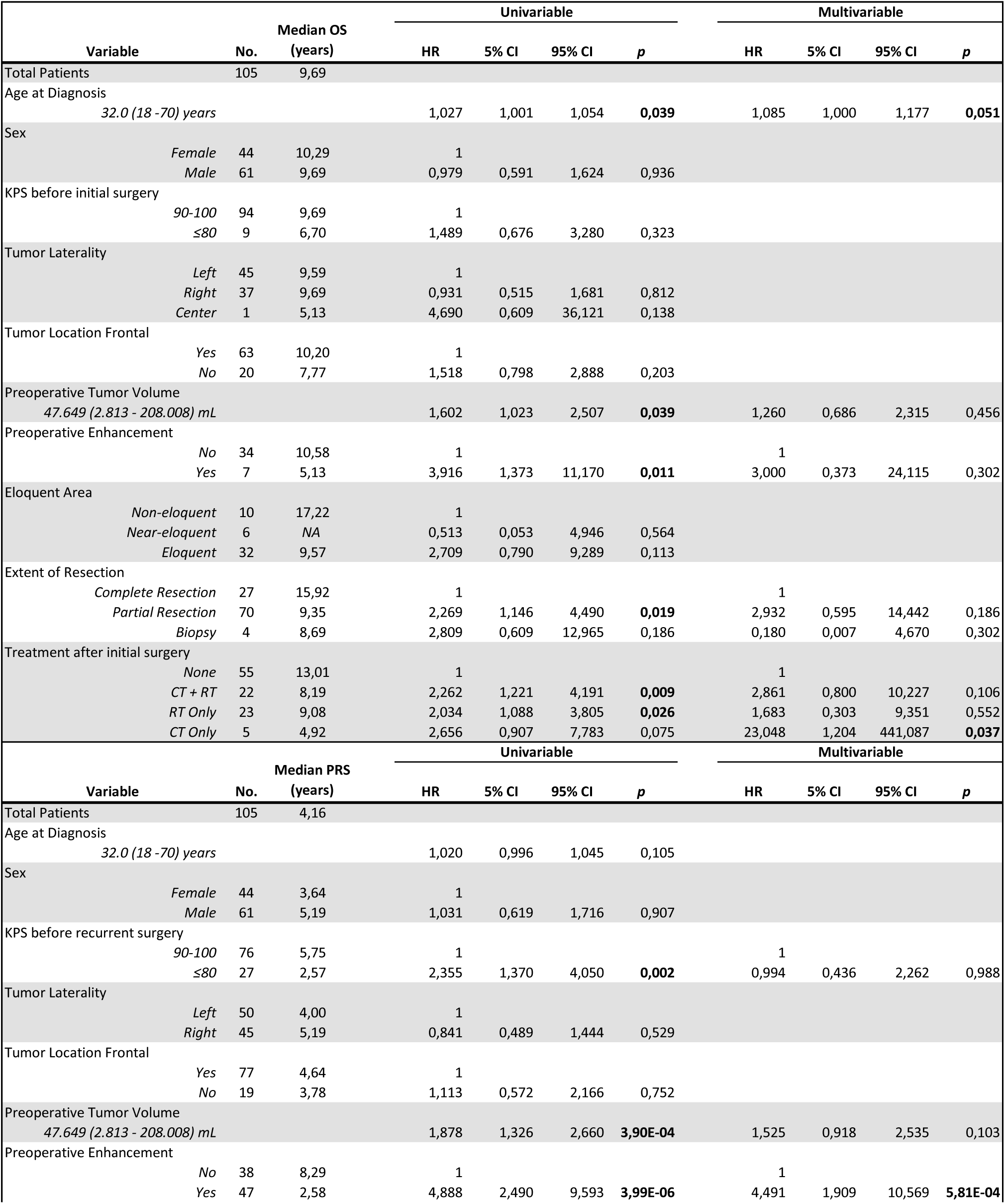
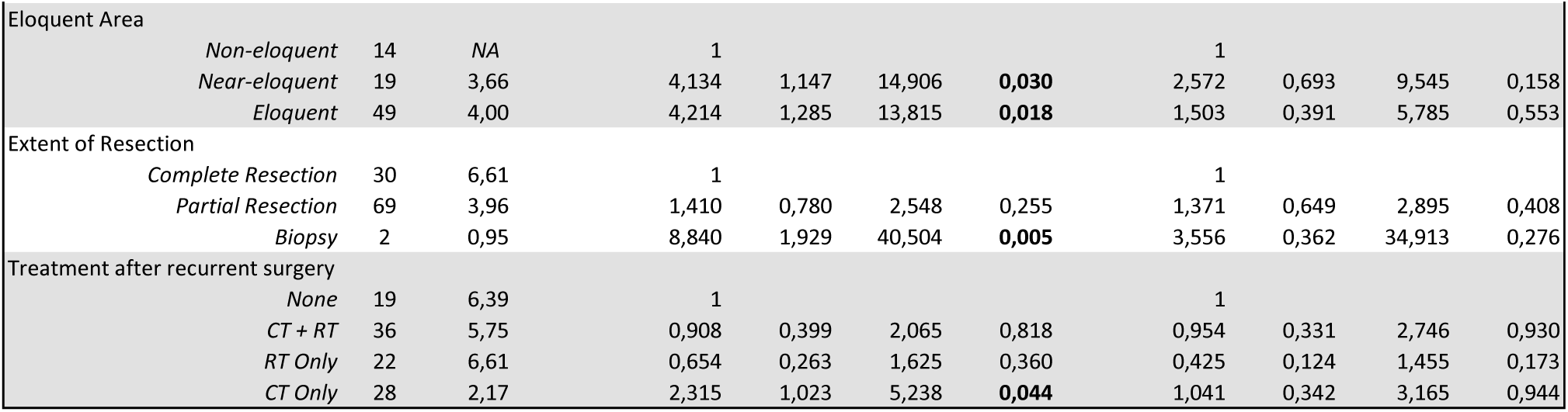
Correlation of clinical parameters with patient survival. Median OS correlated with clinical parameters related to the initial surgery (upper panels) and median PRS correlated with clinical parameters related to the recurrent surgery (lower panels). Univariable and multivariable Cox PH models, p-values *<*0.05 are in bold. OS = overall survival, PRS = post-recurrent survival.

### 3.2 Longitudinal genomic changes in IDH-mutant astrocytomas

To prevent purity bias in our molecular analyses[11], we estimated tumor purity using three independent methods, and found no significant differences between initial and recurrent tumors (**Fig. 1b** and **Supplementary Fig. 1a**).

As glioma evolution is thought to be shaped by intratumoral heterogeneity (ITH), we examined the temporal changes in chromosomal copy-number heterogeneity (CNH) as a measure of ITH[12]. CNH (ranging from 0.004 to 0.274) was significantly higher in recurrent tumors in a paired analysis (**Fig. 1c**), and was negatively associated with PRS (*p =* 3.18*10*^−^*^7^, HR = 1.129[1.078-1.183]). ITH is a result of genomic instability, defined as the tendency of the genome to acquire genomic alterations during cell division. Chromosomal instability (CIN), a subtype of genomic instability that specifically refers to the gain or loss of larger stretches of DNA, was estimated using two independent methods. Firstly, we found that the arm-level aneuploidy score (AS) (ranging from 0 to 39), was significantly higher at recurrence, partially driven by a subset of tumors with a high temporal increase (**Fig. 1d**). High AS was negatively associated with OS in the initial tumors (*p =* 0.005, HR = 1.122[1.036-1.216]), and PRS in the recurrent tumors (*p =* 3.8*10*^−^*^13^, HR = 1.155[1.111-1.2]). Secondly, the total CNV-load (ranging from 0.1 to 1.1 log2(Mb)), was also significantly increased at tumor recurrence (**Fig. 1e**), and associated with poor OS in the initial tumors (*p =* 0.032, HR = 1.383[1.029-1.859]) and PRS in the recurrent tumors (*p =* 3.64*10*^−^*^7^, HR = 2.378[1.703-3.32]). All measures of genomic instability were positively correlated (**Fig. 1f**), with a subset of IDH-mutant astrocytoma that become considerably more unstable over time.

The sample-specific genome-wide copy-number alteration (CNA) patterns were diverse though some chromosomal arms were more frequently affected (**Fig. 1g**). Specifically chr1p12-q23.3, chr3p24.3-p11.1, chr9p24.3-p21.2, chr10q21.3-q26.3, chr14q11.2-q32.33, and chr22q11.21-q13.33 were more commonly affected in recurrent tumors (**Supplementary Fig. 1b**, see also references[13–15]). Loss of chr9p24.3-p21.2 and chr10q21.3-q26.3 (encompassing the *CDKN2A/B* and PTEN locus respectively) in recurrent tumors were negatively associated with PRS (**Supplementary Fig. 1c**). Homozygous deletion (HD) of *CDKN2A/B*is a molecular marker of poor prognosis, introduced in the WHO CNS5 to assign grade 4 to IDH-mutant astrocytomas harboring this deletion. We found that *CDKN2A/B* HD specifically, was associated with high AS in initial and recurrent tumors (*p =* 1.65*10*^−^*^4^ & *p =* 6.08*10*^−^*^10^), and high CNV-load and CNH in recurrent tumors (*p =* 6.96*10*^−^*^8^& *p =* 0.001, Wilcoxon rank sum test).

The top ranking mutated genes were *IDH1* (92,9%, 197/212), *TP53* (74,0%, 157/212) and ATRX (71,2%, 151/212) (**Fig. 1h**). Over time, three patients lost their *IDH1* mutation completely. Five recurrent tumors gained mutations in either*PIK3CA* and PIK3R1, which are most commonly associated with OS in IDH-mutant astrocytomas[16–18]. We did not find any specific genes more frequently mutated in either initial or recurrent tumors in a paired analysis (McNemar’s *χ*^2^ test, FDR *<*0.05, BH-corrected) although the total mutation count was significantly higher in recurrent tumors (*p =* 0.004, Wilcoxon signed-rank test). This increase originated from a small number of recurrences which gained a relatively large number of mutations, as the mean tumor mutational burden (TMB) was almost doubled (3.5 vs 6.8 muts/Mb), while the median was only slightly higher in recurrent tumors (2.7 vs 3.0). High TMB (hypermutation) has been associated with higher tumor grade and shorter OS in IDH-mutant astrocytomap[5,19]. A total 20 tumors of the GLASS-NL cohort were hypermutated (*≥*6 muts/Mb, **Supplementary Fig. 1d**). Although the proportion of hypermutant tumors had doubled at recurrence, this increase was not significant (6 vs 14: *p =* 0.24, Fisher’s exact test). Notable, patients with hypermutant recurrences had significantly shorter PRS, while time-to-recurrence (TTR) did not differ (PRS: *p =* 1.44*10*^−^*^6^, HR = 5.83[2.846-11.94]; TTR: *p =* 0.188, HR = 0.653[0.3469-1.23]). In addition, the hypermutator phenotype was also associated with high CNH in recurrent tumors (*p =* 0.013) and high AS and CNV-load in both initial and recurrent tumors (AS: *p =* 0.04 & *p =* 4.45*10*^−^*^4^; CNV-load: *p =* 0.005 & *p =* 8.48*10*^−^*^5^, Wilcoxon rank sum test).

Previous studies reported that some tumors develop the hypermutator phenotype due to an increase of mutations in mismatch repair (MMR) genes in response to temozolomide (TMZ) chemotherapy[19–21]. Indeed, patients treated with chemotherapy showed higher incidences of the hypermutator phenotype (‘CT + RT’ (6/19) vs ‘None’ (1/49): *p =* 0.009; ‘CT Only’ (4/5) vs ‘None’: *p =* 4.67*10*^−^*^4^; ‘CT Only’ vs ‘RT Only’ (1/23): *p =* 0.007, Pairwise Fisher’s exact test, Bonferroni-corrected), more specifically with TMZ chemotherapy (9/21 vs 2/72; *p =* 1.28*10*^−^*^5^, Fisher’s exact test). Hypermutant status was significantly more abundant in recurrent tumors with mutations in MMR genes (*MSH2, MSH4, MSH5, MSH6, MLH1, MLH3, PMS1* and *PMS2* ; 6/10 vs 6/86 without: *p =* 1.65*10*^−^*^4^, Fisher’s exact test). It is suggested that ionizing radiation induces unique DNA-damage patterns, possibly promoting the acquisition of *CDKN2A/B* HD[22]. *CDKN2A/B* HD was indeed more abundant in recurrent tumors treated with both radio- and chemotherapy, but the difference was not significant in tumors treated with radiotherapy alone (‘CT + RT’ (13/17) vs ‘None’ (8/45): *p =* 1.78*10*^−^*^4^; ‘RT Only’ (8/23) vs ‘None’: *p =* 0.834, Pairwise Fisher’s exact test, Bonferroni-corrected).

### 3.3 Identification of prognostic DNA-methylation signature

The heterozygous mutation in *IDH1/2* is one of the earliest genetic alteration in IDH-mutant gliomas, and due to its indirect inhibitory effect on TET-mediated DNA-demethylation, IDH-mutant gliomas generally exhibit a hypermethylation phenotype[23,24]. We found that over time the genome-wide DNA-methylation levels tend to decrease, as recurrent tumors had, on average, significantly lower DNA-methylation levels compared to initial tumors, in a paired analysis (**Fig. 2a**). In an effort to identify underlying mechanisms of temporal DNA-demethylation, we studied the differentially methylated regions (DMRs) between initial and recurrent tumors. In total, 44,623 DMRs were identified, and on chromosome 6p22.2-p21.2 an enrichment of DMRs could be observed, several of which were also highly significant (**Fig. 2b**). This suggests that in recurrent tumors, DNA-methylation is specifically lifted from this locus to allow for transcription of target genes. Pathway enrichment analysis of the DMRs in promoter regions on 6p22.2-p21.2, revealed strong associations with the nucleosome, DNA packaging, chromatin assembly and chromosome organization (**Fig. 2c**).

**Fig. 2:**
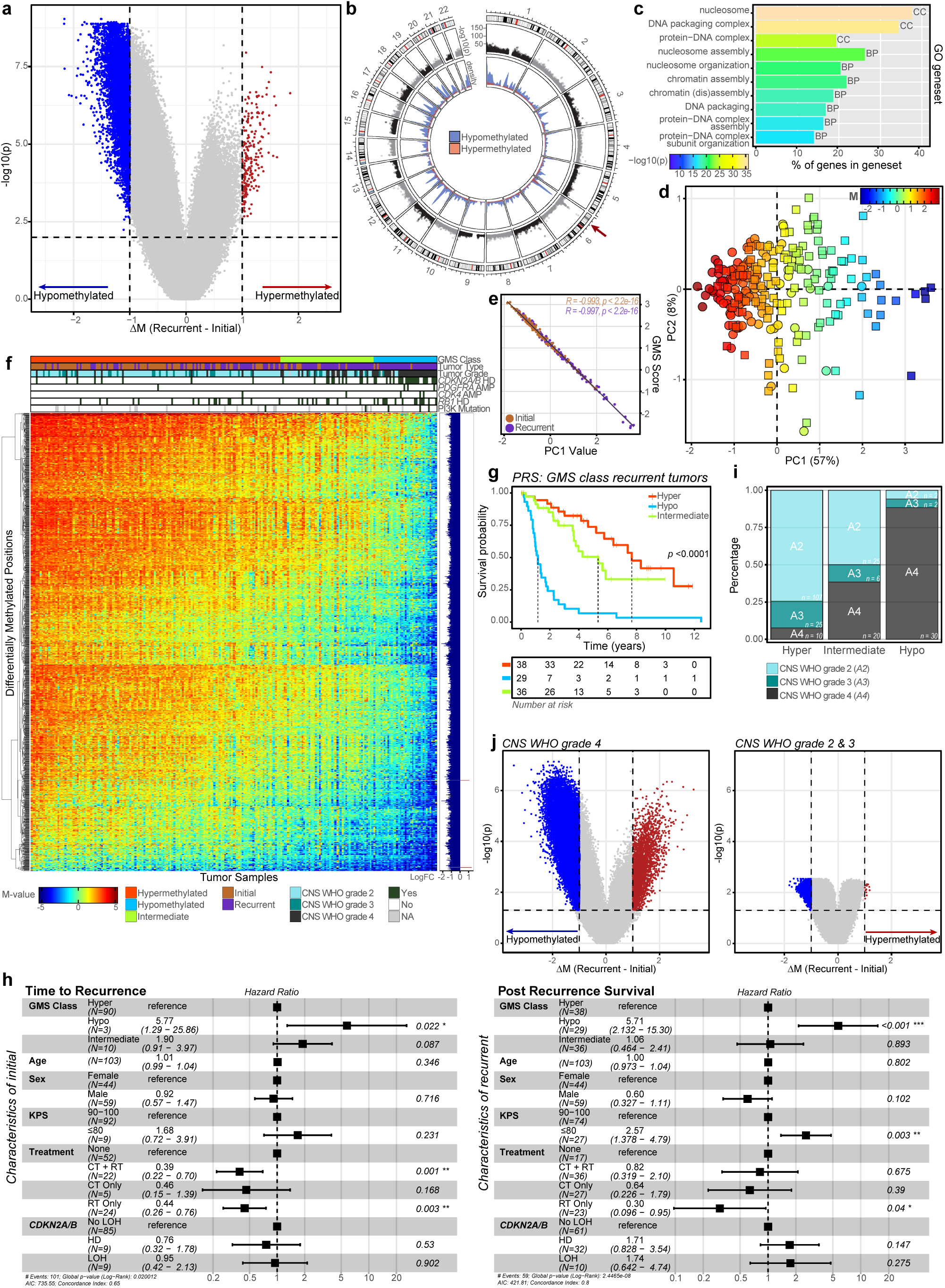
Temporal changes in DNA-methylation and identification of glioma methylation signature. **a**, Volcano plot depicting the probe-level temporal changes in DNA-methylation. Significantly hypermethylated (in red, n = 219) and hypomethylated (in blue, n = 7,483) CpGs (out of 656,517 CpGs in total) between initial and recurrent tumors were identified using a paired Wilcoxon signed-rank test (BH-corrected, FDR *<*0.01 & ΔM *>*1). **b**, Circos plot depicting the 44,623 DMRs identified using a patient-corrected linear regression (BH-corrected, FDR *<*0.05) on their chromosomal location. These regions are characterized by the presence of *≥*2 DMPs per 1,000 nucleotides. The -log10(p-value) per DMR, and the density of hypomethylated (blue) and hypermethylated (red) DMRs are depicted on the outer and inner ring respectively. Enrichment on chromosome 6 is indicated with a red arrow. **c**, GO enrichment analysis of genes overlapping with DMRs in promotor regions on chromosome 6p22.2-p21.2. Top 10 most significant pathways are shown, the -log10(p-values) for overrepresentation of the gene set are FDR adjusted using BH correction. X-axis depicts the percentage of genes from the gene set that were differentially methylated, GO subontologies are indicated at the end of the bar for each gene set. **d**, PCA on DNA-methylation data of initial and recurrent tumor samples, using only the 1,389 highly differentially methylated DMPs from a patient-corrected linear regression (BH-corrected, FDR *<*10*^−^*^9^ & absolute LogFC *>*1). Circles represent initial and squares represent recurrent tumor samples, all shapes are colored based on the median M-value of the selected 1,389 DMPs. **e**, Pearson’s correlation between the median M-value of the selected 1,389 DMPs (GMS score) and PC1 (PCA from **d**) in initial (R = -0.993, *p<*2.2*10*^−^*^16^) and recurrent (R = -0.997, *p<*2.2*10*^−^*^16^) tumor samples separately (with 95% CI). **f**, Heatmap of DMPs from the GMS. Tumor samples are ranked based on the median M-value of the selected 1,389 DMPs (GMS score). Column annotation shows the GMS class, tumor type, CNS WHO grade and presence of molecular markers per sample and row annotation shows the patient-corrected regression coefficient (LogFC). **g**, Kaplan-Meier curve for PRS when stratifying patients on the GMS class of the recurrent tumor. GMS classes were based on their GMS score (‘hyper’: *>*1 M; ‘intermediate’: 0 – 1 M; ‘hypo’: *<*0 M). The hypo GMS class of the recurrent tumors was, as compared to the hyper GMS class, negatively associated with PRS (*p*= 4.1*10*^−^*^9^ HR = 7.037 [3.6723-13.484]). **h**, Multivariate Cox PH models on TTR and PRS with characteristics of initial and recurrent respectively. i, Distribution of CNS WHO grades within the three GMS classes. Cases in the hyper GMS class were predominantly grade 2 (107/142, 75.4%), while grade 4 tumors made up the highest proportion of samples in the hypo GMS class (30/34, 88.2%). **j**, Volcano plots depicting the probe-level temporal changes in DNA-methylation when stratifying patients based on the CNS WHO grade of the recurrent tumor. Differentially methylated CpGs were identified for patients with CNS WHO grade 4 (3041 hyper- vs 33982 hypomethylated CpGs) and CNS WHO grade 2 or 3 (12 hyper- vs 1998 hypomethylated) recurrences using a Wilcoxon signed-rank test (BH-corrected, FDR *<*0.05 & ΔM *>*1). DMR = differentially methylated region, DMP = differentially methylated position, BP = Biological Process, MF = Molecular Function, CC = Cellular Component, PCA = principal component analysis, GMS = glioma methylation signature, TTR = time to recurrence, PRS = post-recurrent survival.

We selected the 1,389 most differentially methylated positions (DMPs) between initial and recurrent tumors. Principal component analysis based on only these DMPs, did not result in two distinct clusters for initial and recurrent tumors, but instead shows a gradient on principal component 1 (PC1), strongly anti-correlating with the median M-value of the 1,389 selected DMPs (**Fig. 2d,e**). This indicates that most of the variance between the tumor samples is explained by the median level of DNA-methylation of the tumor sample. Further, we found that the median M-value of these DMPs in recurrences was strongly associated with OS (*p =* 4.62*10*^−^*^5^, HR = 0.663[0.544-0.808]) and PRS (*p =* 1.26*10*^−^*^7^, HR = 0.607[0.504-0.730]), the higher the value, the better the outcome. We therefore appointed the median M-value of this probe-set as the per-sample ‘glioma methylation signature’ (GMS) score. Of note, our procedure to discover this signature deviates from one based on ‘standard classifiers’ (such as logistic lasso regression) for two reasons. First, the paired as opposed to independent samples setting of this study, and second, our preference for a simple, interpretable score over a more complex one, as usually rendered by such classifiers.

Ranking all tumor samples on the GMS score shows that the initial tumors mostly cluster towards the high-methylation end of this spectrum, while recurrent tumors are more abundant at the low-methylation end (**Fig. 2f**). Furthermore, the smooth gradient suggests that the observed temporal DNA-demethylation is likely a secondary event. Therefore, in an effort to unravel what drives temporal DNA-demethylation, we examined the association of our signature, with known molecular markers for progression (*CDKN2A/B* HD, *CDK4* amplification, PDGFRA amplification, RB1 HD and/or PI3K mutations)18. We found that low GMS score was associated with the presence of any of these markers (initial: 19/103, *p =* 0.003; recurrent: 49/103 *p =* 2.62*10*^−^*^6^), even when eliminating cases with only *CDKN2A/B* HD (initial: 12/103, *p =* 0.002; recurrent: 29/103 *p =* 0.001, Wilcoxon rank sum test), suggesting that genetic changes underlie the observed temporal DNA-demethylation.

The GMS score was linearly associated with patient survival, but for the ease of use in daily clinical practice we split our samples into three GMS classes (‘hyper’: *>*1 M; ‘intermediate’: 0 – 1 M; ‘hypo’: *<*0 M). The hypo GMS class in recurrent tumors was negatively associated with PRS (**Fig. 2g**), and its prognostic value was independent of known clinically prognostic factors (age, sex and performance), treatment (radio- and/or chemotherapy) and molecular markers (*CDKN2A/B* HD), in a multivariable analysis (**Fig. 2h**). Considering that the hypo GMS class outperforms *CDKN2A/B* HD, this indicates that our signature should be regarded as a grading aid for IDH-mutant astrocytomas.

Continuing, we also found a correlation between the GMS classes and tumor grade, as tumors of the hyper GMS class were predominantly grade 2, and those of the hypo GMS class were mostly grade 4 tumors (**Fig. 2i**). These grade differences account for much of the observed temporal DNA-demethylation, since tumors that recur as grade 4 show significantly decreased genome-wide DNA-methylation levels, while the temporal changes in DNA-methylation are limited in tumors that recur as grade 2 or 3 (**Fig. 2j**). Increased incidences of high-grade tumors at recurrence therefore seem to be associated with DNA-demethylation.

To validate whether the GMS was also prognostic in newly diagnosed IDH-mutant astrocytoma patients, we used DNA-methylation data from 432 patients included in the randomized phase III CATNON clinical trial[17,25]. Also in this dataset, the tumor samples could be gradually separated based on the per-sample GMS score (**Fig. 3a**), which was also strongly associated with the presence of malignancy markers (n = 103/432, *p =* 9.45*10*^−^*^16^), even when eliminating cases with only *CDKN2A/B* HD (n = 67/432, *p =* 2.595*10*^−^*^10^, Wilcoxon rank sum test) (**Fig. 3b**). The lower GMS classes were negatively associated with OS and progression-free survival (PFS) (**Fig. 3c**), and in a multivariable analysis, poor survival of the hypo GMS class was independent of known clinically prognostic factors, additional treatment and *CDKN2A/B* HD (**Fig. 3d**). This validates, in an independent dataset, the prognostic value of our signature, further strengthening the notion that it should be appreciated as grading aid for IDH-mutant astrocytomas.

**Fig. 3:**
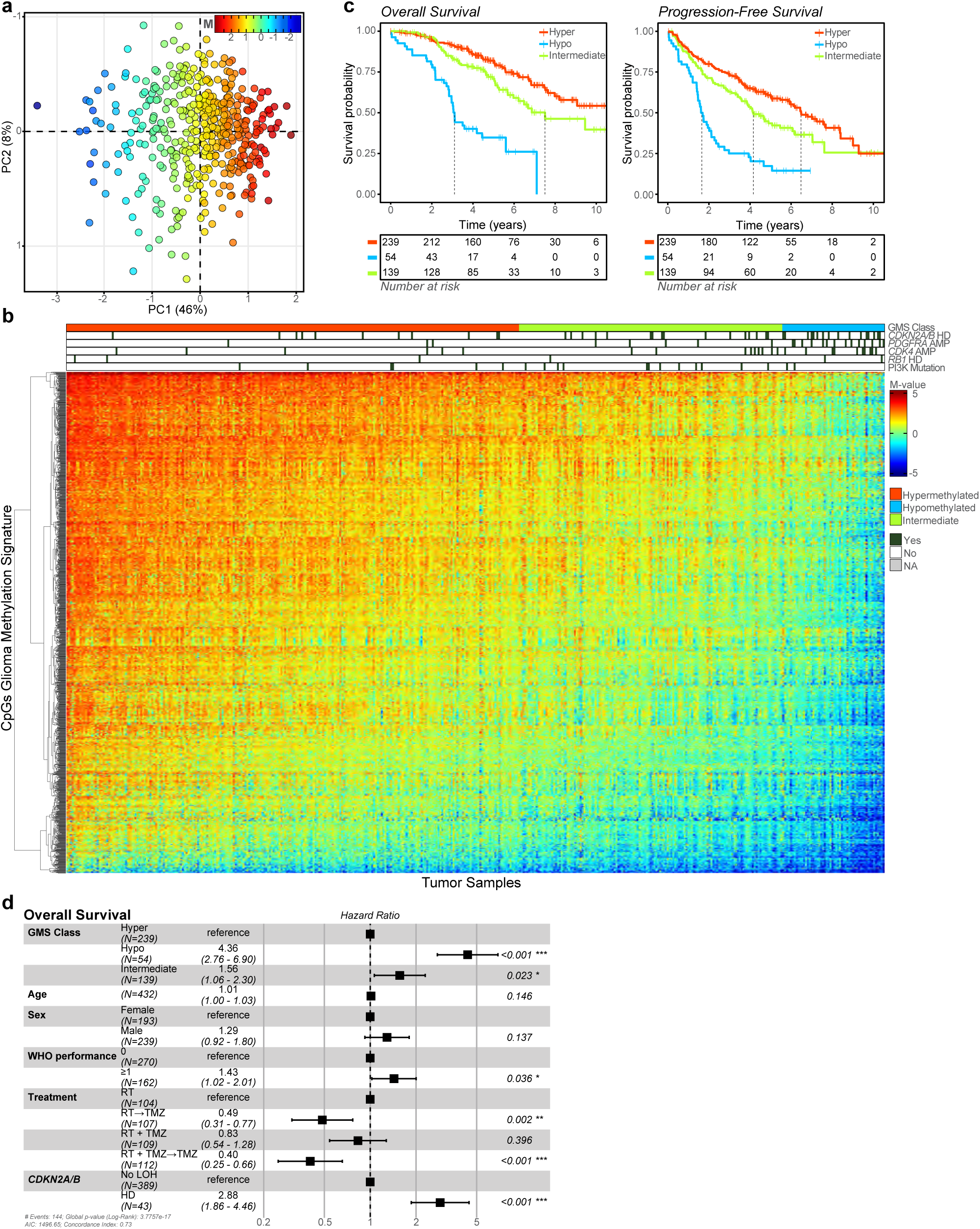
Validation of the ‘glioma methylation signature’. **a**, PCA on DNA-methylation data of all tumor samples from the CATNON cohort, using only the DMPs from the GMS. Shapes are colored based on the median M-value of the signature CpGs. **b**, Heatmap of DMPs from the GMS. Tumor samples are ranked based on the GMS score. Column annotation shows the GMS class and presence of molecular markers per sample. **c**, Kaplan-Meier curves for OS and PFS when stratifying patients on the GMS class. The lower GMS classes were negatively associated with OS (‘hypo’: p = 5.53*10*^−^*^14^, HR = 5.109 [3.339-7.815]; ‘intermediate’: p = 0.012, HR = 1.625 [1.111-2.376]), but also with PFS (‘hypo’: p = 1.64*10*^−^*^11^, HR = 3.438 [2.400-4.924]; ‘intermediate’: p = 0.013, HR = 1.452 [1.082-1.947]). **d**, Multivariate Cox PH model on OS. PCA = principal component analysis, DMPs = differentially methylated positions, GMS = glioma methylation signature, OS = overall survival, PFS = progression-free survival.

### 3.4 Cell cycling strongly upregulated over time in IDH-mutant astrocytomas

To investigate the temporal transcriptomic changes, we performed differential expression analysis between initial and recurrent tumor samples and identified 604 differentially expressed genes (DEGs). A subset of upregulated DEGs were concentrated on chromosome 6p22.2-21.33, overlapping the region enriched with hypomethylated DMRs (**Fig. 4a**). The genes on this locus are mainly histone genes, encoding for proteins involved in the formation of histone-complexes and therefore nucleosome formation and DNA-packaging. Co-expression clustering analysis of all 604 DEGs revealed four clusters, three representing up- and one representing downregulated genes at recurrence (C1-C4) (**Fig. 4b**). To uncover the processes involved in these transcriptional changes, we investigated the associations and cell type specific expression of these gene clusters. We performed single-nucleus RNA-sequencing (snRNA) on flash-frozen material of three matching tumor pairs of the GLASS-NL cohort, and complemented these data with single-cell RNA-sequencing (scRNA) data pooled from three independent IDH-mutant astrocytoma datasets[26–28]. Of note, the recurrent tumors of all three GLASS-NL patients were selected to be of grade 4.

**Fig. 4:**
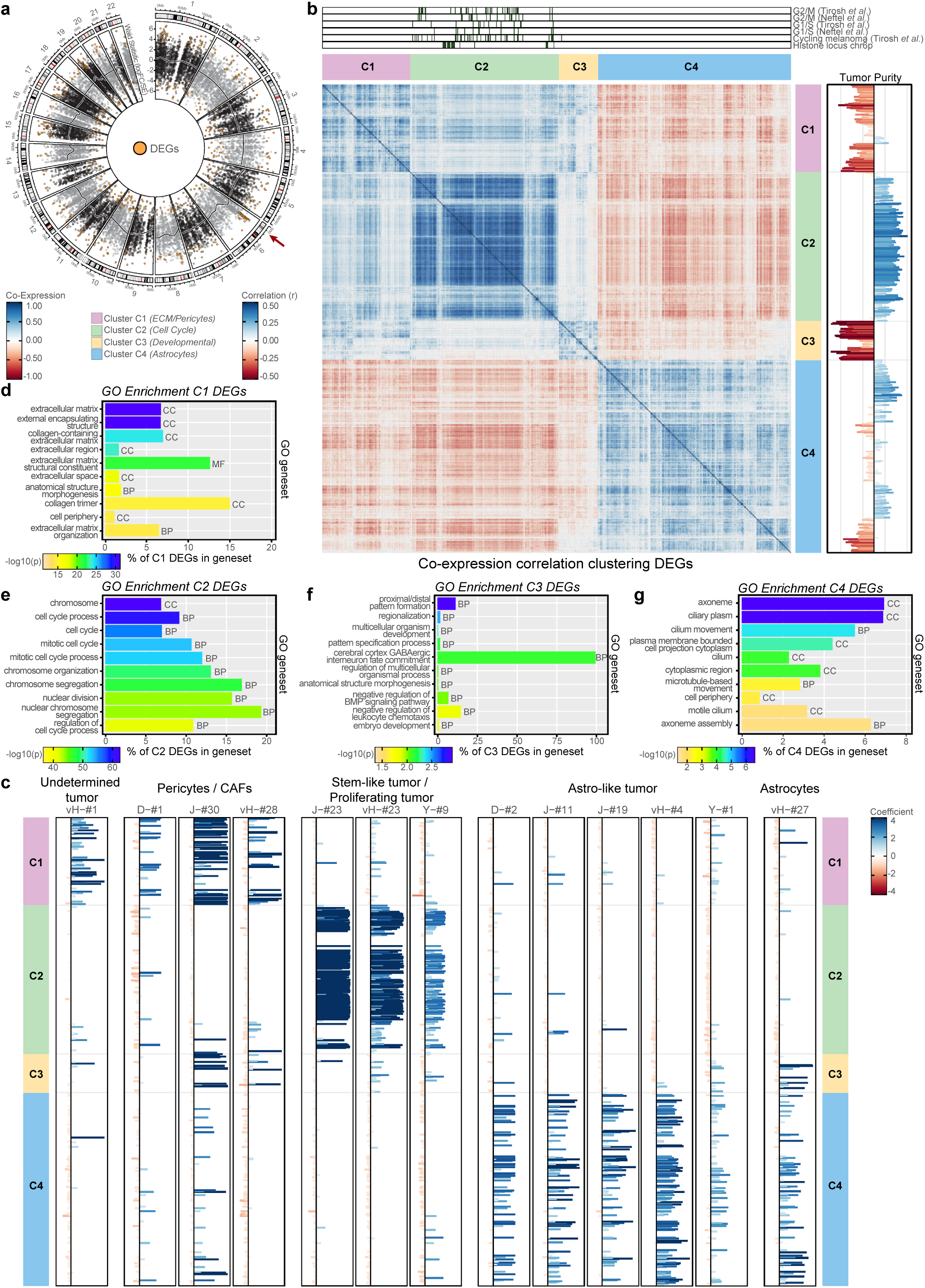
Temporal changes in RNA expression and identification of co-expression clusters. **a**, Circos plot depicting the assessed genes on their chromosomal location. The Wald statistic (LogFC/SE) of each gene is plotted, for each chromosome the smoothed conditional means (formula = y ’s(x, bs = cs), method = REML) is drawn. Differentially expressed genes between initial and recurrent tumors were identified using DESeq2 (BH-corrected, FDR *<*0.01 & absolute LogFC *>*0.75). The 604 DEGs (355 up- and 249 downregulated) are colored in orange. Enrichment on chromosome 6 is indicated with a red arrow. **b**, Recursive correlation plot depicting the co-expression correlation clustering of the 604 DEGs. Four clusters are identified and are represented by colors (C1: pink, C2: green, C3: yellow, C4: blue). Column annotations (top) indicate cell cycle genes from literature and genes from the chr6p histone locus. Row annotation shows the correlation (r) to tumor purity. **c**, Association of cell type specific annotations from the scRNA/snRNA datasets to the genes from the expression clusters (C1-C4). Cell type specific clusters from the datasets with the highest average expression of the 604 DEGs are visualized. The annotation of cell types was diverse among datasets, in the Johnson dataset[28] cycling tumor cells were annotated as ‘proliferating stem-like’ and ‘stem-like’. **d-g**, GO enrichment analysis of genes in expression clusters C1-C4. Top 10 most significant pathways are shown, -log10(p-values) for overrepresentation of the gene set are FDR adjusted using BH correction. X-axis depicts the percentage of genes from the gene set that were present in the cluster, GO subontologies are indicated at the end of the bar for each gene set. DEGs = differentially expressed genes, BP = Biological Process, MF = Molecular Function, CC = Cellular Component.

The upregulated cluster C1 is mainly characterized by collagens and the expression of its genes are inversely correlated with tumor purity (**Fig. 4b**). These genes are predominantly expressed by pericytes and pathway enrichment analysis revealed associations with the extracellular matrix (**Fig. 4c,d**). This cluster showed a high overlap with an expression cluster identified in IDH-wildtype glioblastomas, whose genes were also temporally upregulated and expressed by pericytes11. This overlap suggests that both IDH-mutant astrocytomas and IDH-wildtype glioblastomas adopt similar mechanisms to modify the extracellular matrix to enhance tumor malignancy. Cluster C2 encompasses all upregulated genes from the chromosome 6p22.2-21.33 histone locus, and is further enriched with genes that are used to define cycling cells, including*TOP2A*[29,30] (see top-annotation in **Fig. 4b**). Hence, pathway enrichment analysis revealed associations with cell cycle processes and chromosome organization (**Fig. 4e**). Its expression originates predominantly from tumor cells as the expression of genes from C2 strongly correlated with tumor purity, and these genes were mainly expressed by cycling tumor cells (**Fig. 4c**). Genes from C3 showed less inter-cluster correlation and no enriched cell types were found, pathway enrichment analysis revealed mainly associations with development associated gene-sets, albeit not very strongly (**Fig. 4f**). The downregulated genes from cluster C4, contains markers for mature astrocytes including *GABRG1* [31], the few enriched pathways found were all associated with cilia (**Fig. 4g**). These genes are predominantly expressed by astrocytes and astrocyte-like tumor cells (**Fig. 4c**), and the observed temporal change in expression may originate from either. In summary, we find that the fraction of astrocyte-like tumor cells is reduced over time, while the fraction of dividing tumor cells increases, resulting in the observed differences in expression.

### 3.5 Increased cell cycling originates from high-grade recurrences

To validate the temporal differences in expression of gene clusters C1-C4, we integrated the snRNA data of our three IDH-mutant tumor pairs (**Fig. 5a**). The integrated dataset showed clear overlap of cells from all tumor samples and coverage of all major cell types expected in the tumor microenvironment (**Supplementary Fig. 2a,b**). All gene clusters showed distinct (temporal) expression patterns (**Fig. 5b**). More specifically, genes from C2 showed enrichment in the proliferating tumor cell fraction of recurrent tumors, C3 genes were mainly expressed by recurrent tumor cells that were not differentiated into oligo-like or astro-like tumor cell states, while C4 expression was enriched in the astro-like tumor cell fraction of initial tumors. The combined temporal upregulation of C3, and downregulation of C4 is indicative of tumor cell de-differentiation over time. Furthermore, the temporal changes in clusters C2 and C4 were conserved in all three patients with high grade recurrences (**Fig. 5c** and **Supplementary Fig. 2c**).

**Fig. 5:**
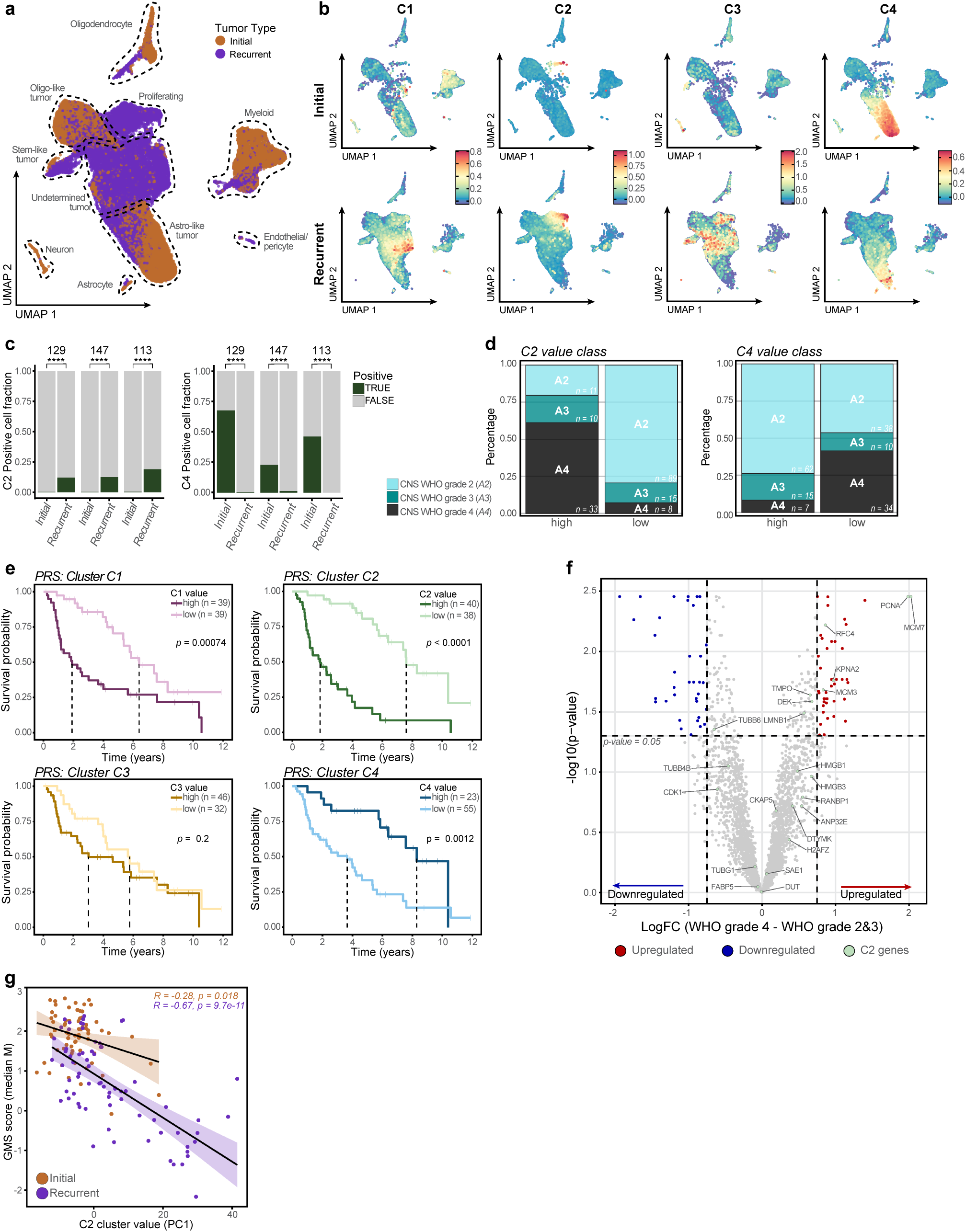
Single-cell and single-nucleus RNA sequencing validation of expression clusters. **a**, UMAP projection of integrated single nuclei of initial and recurrent tumors from three IDH-mutant astrocytoma patients. Different cell types are annotated within the plot (see also **Supplementary Fig. 2a,b**). **b**, UMAP projection of single nuclei of initial (top panels) and recurrent tumors (lower panels) for expression clusters C1-C4. Nuclei are colored by the enrichment score calculated for the concerning expression clusters. **c**, Barchart of the positive cell fractions of expression clusters C2 and C4 of each individual patient (see also **Supplementary**Fig. 2c). Difference in positive cell fraction between initial and recurrent tumor sample in each patient is calculated using a Pearson’s *χ*^2^ test. In all three patients there was a highly significant co-directional difference in positive tumor cell fraction between initial and recurrent tumors for both expression clusters (p-value was in all cases *<*2.2*10*^−^*^16^). * p-value *<*0.05, ** p-value *<*0.01, *** p-value *<*0.001, **** p-value *<*0.0001. **d**, Distributions of CNS WHO grades within the expression cluster value classes (‘high’: PC1 *>*0; ‘low’: PC1 *<*0) of C2 and C4 (graphs of C1 and C3 are found in **Supplementary Fig. 2d**). C2 ‘high’ cases were predominantly grade 4 (33/54, 61.1%) and C2 ‘low’ cases grade 2 (89/112, 79.5%). Contrary, C4 ‘high’ cases were predominantly grade 2 (62/84, 73.8%) and 41.5% (34/82) of the C4 ‘low’ cases were grade 4. **e**, Kaplan-Meier curves for PRS of expression cluster C1-C4. Patients were stratified based on the cluster value classes (‘high’: PC1 *>*0; ‘low’: PC1 *<*0) of the recurrent tumor. The continuous cluster values of the recurrent tumor samples showed similar prognostic value in Cox PH models on PRS (C1: *p*= 7.47*10*^−^*^7^, HR = 1.063[1.037-1.089], C2: *p*= 1.96*10*^−^*^14^, HR = 1.086[1.063-1.109], C3: *p*= 0.344, HR = 1.031[0.968-1.098], C4: *p*= 1.47*10*^−^*^7^, HR = 0.910[0.879-0.943]). **f**, Volcano plot depicting the DEPs between CNS WHO grade 2&3 and CNS WHO grade 4. Significantly upregulated (in red, n = 44) and downregulated (in blue, n = 36) genes were identified using an unpaired linear regression model (BH-corrected, FDR *<*0.05, absolute LogFC *>*0.75). Genes from the C2 cell cycle expression cluster are annotated in green. **g**, Pearson’s correlation between the GMS score and the C2 cluster value in initial (R = -0.28, *p*= 0.018) and recurrent (R = -0.67, *p* = 9.7*10*^−^*^11^) tumor samples separately (with 95% CI). PRS = post-recurrent survival, DEPs = differentially expressed proteins, GMS = glioma methylation signature.

**Fig. 6:**
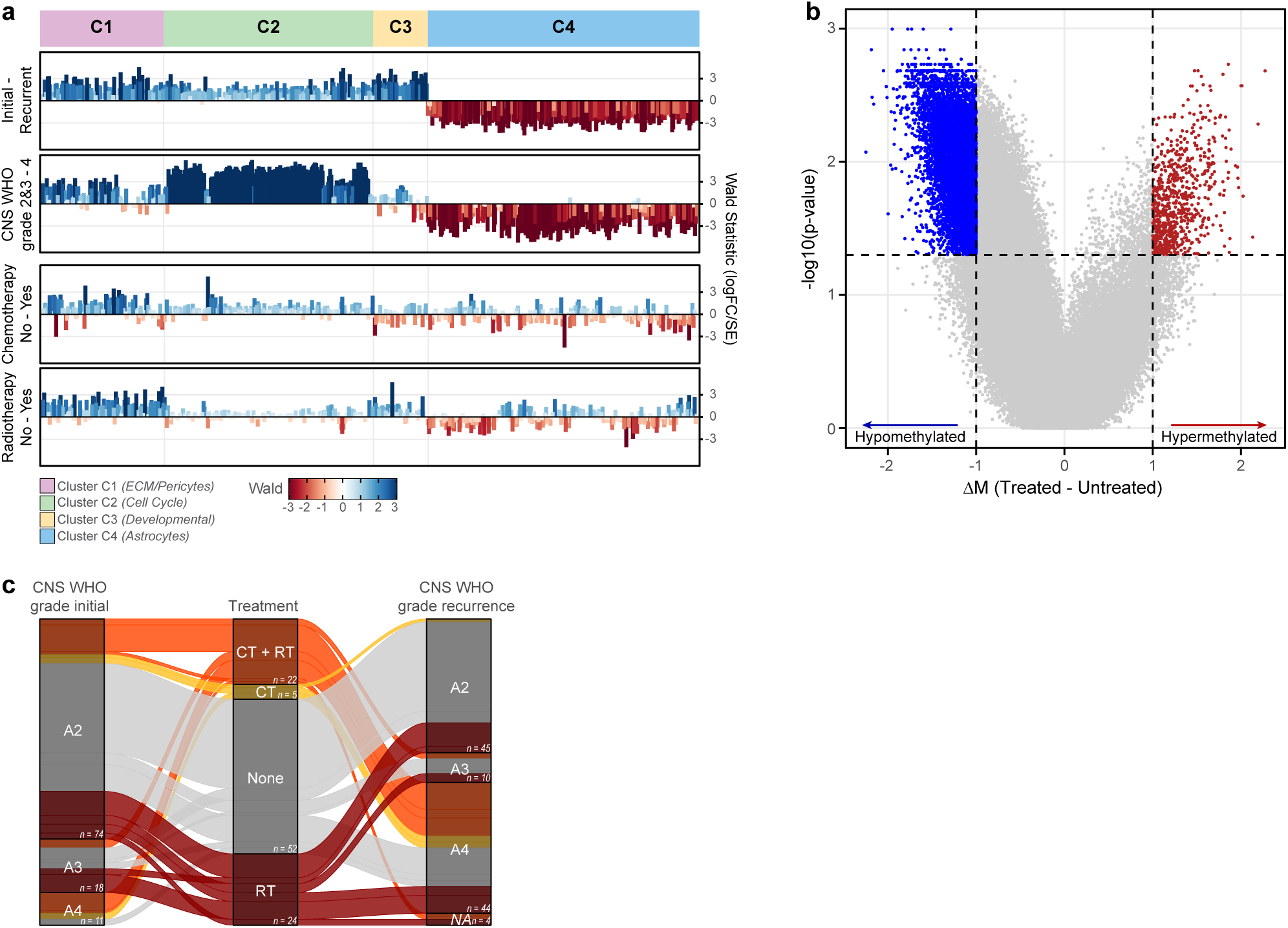
Effect of chemo- and radiotherapy on RNA expression and DNA-methylation. **a**, Annotation to the recursive correlation (as visualized in **Fig. 4b**) of the multivariate differential expression analysis. The contribution of confounding factors (from top to bottom: initial-recurrent, WHO grade 2&3 - 4, treated with chemotherapy, and treated with radiotherapy) to the expression of genes within the expression clusters C1-C4 is measured by the Wald statistic (LogFC/SE). **b**, Volcano plot depicting the probe-level differences in DNA-methylation of recurrent tumors with (n = 51) and without (n = 52) prior chemo- and/or radiotherapy treatment. Differentially hypermethylated (in red, n = 738) and hypomethylated (in blue, n = 10,527) CpGs between treated and untreated recurrent tumors were identified using a Wilcoxon rank sum test (BH-corrected, FDR *<*0.05 & ΔM *>*1). **c**, Sankeyplot visualizing the CNS WHO grade of initial and recurrent tumors in relation to additional treatment. Treatment interventions (limited to radio- and/or chemotherapy) between initial and recurrent surgery were aggregated into the projected treatment groups. Most of the tumors that recurred as CNS WHO grade 4 have been previously treated (32/44, 73%).

Since all three snRNA tumor pairs displayed malignant transformation at recurrence, we evaluated the effect of tumor grade on the expression clusters, in a multivariate model including other covariates (visualized in **Fig. 6**). The effect of tumor grading on clusters C2 and C4 was stronger than the temporal (initial-recurrent) effect. This indicates that the temporal upregulation of C2 and downregulation of C4 genes arise from the increased fraction of high-grade tumors at recurrence. We condensed the gene expression for each cluster into a per-sample cluster value, and split the samples into ‘high’ and ‘low’ expression-classes, and found a correlation between the expression classes of C2 and C4, and tumor grade (**Fig. 5d** and **Supplementary Fig. 2d**). Furthermore, we found that high expression of C1 and C2, and low expression of C4 in recurrences, was negatively associated with survival (**Fig. 5e**), similar prognostic value was observed when using the continuous cluster values. The expression of C3 did not associate with survival.

The upregulation of cell cycling associated genes is also reflected in the protein expression data. Unpaired differential protein expression analysis indicated 80 proteins to be significantly differentially expressed between low- and high-grade tumor samples (**Fig. 5f**). Many of the upregulated proteins in high-grades are involved in cell cycling, including PCNA and MCM7[32,33], further supporting the notion that cell cycle is elevated in tumors that recur as CNS WHO grade 4 IDH-mutant astrocytomas. There were no differentially expressed proteins between initial and recurrent tumors identified, though this is likely related to the low number of available matched tumor pairs (n = 18).

Because the per-sample DNA-methylation signature score decrease and the upregulation of cell cycle genes were both associated with high-grade recurrences and poor survival, we tested the correlation between these two markers. We found that the C2 cluster value and the GMS score were inversely correlated, indicating that the upregulation of cell cycling is associated with temporal DNA-demethylation, especially in recurrent IDH-mutant astrocytomas (**Fig. 5g**).

### 3.6 Treatment does not induce cell cycling and hypomethylation

Using the above described multivariate model, we evaluated the effect of additional treatment on the gene expression clusters (**Fig. 6a**). While the treatment coefficients are higher for the C1 cluster than for the remaining clusters C2-C4, these coefficients do not exceed those of the non-treatment associated covariates, nor are the multivariate adjusted p-values significant. Treatment is therefore not the driving factor behind the observed temporal transcriptomic changes.

Likewise, we examined whether treatment has a confounding effect on DNA-methylation, by using a mixed-effects model, and aimed to identify DMPs between treated and untreated patients. However, only 13 out of 656,517 CpGs assessed were differentially methylated between patients treated with both radio- and chemotherapy and patients that did not receive any treatment between surgical interventions (FDR *<*0.05, patient-corrected linear regression, BH-corrected), using other stratification methods did not yield more DMPs. The effect of treatment on temporal changes in DNA-methylation does therefore not induce the observed temporal DNA-demethylation in this cohort. We do observe global hypomethylation in recurrent tumors of treated patients compared to those of patients that have not been treated prior to recurrent surgery (**Fig. 6b**). However, our analysis shows that this effect is not induced by treatment itself, but originates from high-grade recurrences which are more abundant in the group of treated patients (**Fig. 6c**).

Together, we conclude that the administration of treatment (i.e. radio- and/or chemotherapy) does not induce the temporal molecular changes observed in this study, and therefore does not promote the observed characteristics associated with malignant transformation of the tumor, in this cohort.

## 4 Discussion

We examined the genomic alterations in matched initial and recurrent IDH-mutant astrocytomas and found that genomic instability increases over time. Loss of large chromosomal regions of chr3p, chr9p, chr10q, chr14q and chr22q were more frequently observed at recurrence and most have been previously reported in IDH-mutant gliomas, and in some cases linked to malignant transformation[13–15]. While the overall number of mutations increased over time, no recurrent-specific gene mutations were found. Of the 212 sequenced tumor samples, 20 were hypermutant, which is less than in other IDH-mutant astrocytoma cohorts7,19. Nevertheless, the hyper-mutant phenotype was more abundant in recurrent tumors of patients that were treated with temozolomide prior to the surgical intervention. In line with previous findings, we observed that genome-wide DNA-methylation decreased over time[13,24]. Changes in tumor grade account for much of this decrease as DNA-demethylation is limited in patients with low-grade, and unambiguous in those with high-grade recurrences. We were able to capture this temporal change in a ‘glioma methylation signature’ (GMS). Tumors with the lowest GMS score more often harbored alterations in known malignancy markers, indicating that it is linked to malignant transformation. Further, our data showed that the GMS was strongly associated with survival, independent of known clinically prognostic factors, treatment regimen and *CDKN2A/B* HD. We validated these findings in a set of newly diagnosed IDH-mutant anaplastic astrocytoma patients[17,34]. The association with malignancy markers and survival indicates that our signature can be used as a molecular tool for tumor grading. This is important because the current grading of IDH-mutant astrocytomas is lacking prognostic power[35], and the difference between CNS WHO grade 2 and 3 is currently ill defined[36–38]. Recent findings from the INDIGO trial underline the importance of accurate grading, as patients with grade 2 IDH-mutant glioma showed strong clinical benefit from *IDH1/2* inhibitor vorasidenib[39], while patients with contrast-enhancing glioma did not respond[40]. A potential added value of our signature is that only a DNA-methylation array is needed to determine the signature score, which are increasingly being used for diagnostic purposes in glial tumors[6,41,42]. In addition to providing an objective means to improve grading of IDH-mutant astrocytomas, we also explain the molecular mechanism underlying tumor malignancy. Focal demethylation of a region on chromosome 6p strongly associated with DNA-packaging and the nucleosome, and the upregulated histone genes on this locus were co-expressed with known cell cycling markers. The increased expression of cell cycle genes predominantly originated from tumor cells. Paired single-nucleus RNA-sequencing analysis further showed that expression of cell cycle genes arises from proliferating tumor cells of recurrent tumors that displayed malignant transformation. The bulk RNA-sequencing data indicated that the upregulation of cell cycling is caused by the increased incidence of high-grade tumors at recurrence. This was supported by the observation that many of the upregulated proteins in high- grade tumors are involved in cell cycling, including PCNA (proliferating cell nuclear antigen) that acts as a DNA clamp and is essential for DNA replication[43]. The upregulation of cell cycle and global DNA-demethylation both seem to be driven by malignant transformation of the recurrent tumors. We also observed that our DNA- methylation signature and cell cycling are correlated, which suggests that they are connected on a biological level. Since DNA-demethylation is most likely a secondary event, we hypothesize that increased cell cycling results in DNA-demethylation. IDH-mutant gliomas are known to display genome-wide DNA-hypermethylation which is most likely due to the competitive inhibition of TET-induced DNA-demethylation[44,45]. During DNA- replication, DNA-methylation is maintained by DNMT1, which is recruited to the replication fork by binding to PCNA[46,47]. It was previously demonstrated that, in highly replicating cells, the maintenance of DNA- methylation cannot keep up with the replication rate, resulting in so called passive DNA-demethylation[48]. We consider this to be an attractive explanation for the observed temporal DNA-demethylation in IDH-mutant astrocytomas in regard to increased cell cycling, but further elucidation of the actual mechanisms underlying this phenomenon are needed. In summary, we studied the molecular changes in IDH-mutant astrocytoma over time. We found that recurrent tumors become genetically more instable, cell cycling is upregulated in high-grade recurrences, which most likely results in passive DNA-demethylation. We propose a signature of 1,389 probes, with strong prognostic power, and a biological rationale, which can be easily implemented in current diagnostics to improve grading of IDH-mutant astrocytomas. Limitations of the study Due to the inclusion criteria of this study, patients included in this cohort are often younger and performing better than the general population of adult IDH-mutant astrocytoma patients. Also, as samples included in this study were obtained in a period of time spanning multiple decades, treatment of patients in this cohort was not homogeneous. Finally, as this study was conducted retrospectively, the (potential) clinical implications should be considered with caution.

## 5 Acknowledgements

This project was funded by the Dutch Cancer Society (KWF Kankerbestrijding), KWF project number 11026/2017-1. The authors acknowledge the contribution of the PALGA foundation in the Netherlands, and the proteomics core facility in Zurich (FGCZ). The CATNON Study was funded by Merck, Sharp & Dohme (MSD) formerly Schering Plough by an educational grant, and by the provision of temozolomide.

## 6 Methods

### 6.1 Patients and sample collection GLASS-NL cohort

In this study we included adult patients that were diagnosed with a histologically proven astrocytoma grade II or grade III at presentation according to the WHO 2016 guidelines49. Histopathological diagnosis and grade were reassessed by a dedicated neuropathologist and reclassified based on molecular features included in the WHO 2021 guidelines[2]. Only patients that underwent surgery at least twice separated by 6 months, and harboring an *IDH1/2* mutation at presentation, were eligible for inclusion. Tumor samples of the initial tumor and its paired recurrences were collected in multiple medical centers in the Netherlands treating patients with gliomas (Erasmus MC Cancer Institute, Rotterdam; Amsterdam UMC Location VUmc, Amsterdam; and UMC Utrecht, Utrecht). For each molecular profiling method, two time-separated tumor samples per patient were selected, composing a matched tumor pair (termed initial and recurrence) for further analysis. In all cases, data from the last available recurrent tumor was selected unless there were any type of quality issues. Surgical interventions on residual tumor or necrotic tissue were excluded for all further analysis. For 105 patients, sufficient material could be collected of both the initial and at least one recurrent tumor, for subsequent genomic, epigenomic, transcriptomic and proteomic analyses. Although there is a high level of overlap in the tumor samples used for all molecular methods, in some cases we were forced to select different recurrences for the molecular analyses in this manuscript. Clinical data was collected by local collaborators using a predefined database, including detailed survival and treatment data. For several analyses, all additional treatments (i.e. radio- and/or chemotherapy) received within a specific time window (adjuvant and at progression), were aggregated into treatment groups (‘CT + RT’, ‘RT Only’ & ‘CT Only’). Treatment after initial surgery comprises all treatments received between the surgical intervention on the initial tumor and that of the selected recurrent tumor, and treatment after recurrent surgery comprises all treatments after surgical intervention of the selected recurrent tumor.

The design of the study was approved of by the institutional review board of Erasmus MC (MEC-2019-0288) and the institutional review board of Amsterdam UMC (MEC-VUMC-2019.085), and the study was conducted according to institutional and national regulations.

### 6.2 DNA and RNA extraction

DNA and RNA from all formalin-fixed-paraffin-embedded (FFPE) tissue blocks, were centrally isolated at Erasmus MC. Tissue areas with high percentage of neoplastic cells (preferably *>*70%, but at least 50%) were manually macro-dissected from 10µm tissue sections. For DNA isolations, the QIAamp DNA FFPE Tissue Kit (Qiagen) was used according to the manufacturer’s instructions with an added overnight proteinase K digestion. For RNA isolations, the RNeasy FFPE Kit (Qiagen) was used according to the manufacturer’s instructions. DNA and RNA concentrations were measured using the Qubit 3.0 Fluorometer according to manufacturer’s protocol (Life Technologies).

### 6.3 IDH mutation and 1p/19q status

*IDH1/2* mutation status of multiple cases was previously determined for diagnostic purposes, by next-generation sequencing or immunohistochemistry of IDH1 R132H. When IDH status was not previously determined, IHC staining of IDH1 R132H was performed on 4-5µm tissue slides of the initial tumor sample, and visually assessed by a dedicated neuropathologist. *IDH1/2* mutation status of cases with negative IHC results were further assessed by sanger sequencing of PCR products of both *IDH1* and *IDH2* regions, generated from genomic DNA. 1p/19q status was previously determined for diagnostic purposes by FISH, MLPA, or copy-number analysis. When 1p/19q status was not previously determined, copy-number profiles extracted from methylation profiling were evaluated for combined deletion of chromosome 1p and 19q.

### 6.4 MRI Features

For all available tumor samples, a pre-operative MR imaging session was selected if available. The scans were scored for location, eloquence and additional features according to the Visually Accessible Rembrandt Images (VASARI) annotations[50].

Tumor segmentations were made if the following sequences were available: T2-weighted (T2w), T2-weighted FLAIR (T2w-FLAIR), and pre- and a post-contrast T1-weighted (T1w / T1w+c) sequences. Scans were excluded if they had severe artifacts such that accurate delineation was not possible. If multiple pre-operative scans were available with sufficient imaging, the latest scan was used. Tumor segmentations were produced automatically using HD-GLIO, and corrected semi-automatically using ITK-SNAP, if needed[51,52]. Two regions of interest were segmented: the area of contrast-enhancing tumor (CET) and non-enhancing hyper-intensities on T2w imaging (NET). The union of both (CET+NET) is considered the whole tumor (WT) volume, necrotic areas were not included. Suspected treatment-induced abnormalities were only excluded if they appeared distant to the original and recurrent tumor lesion and no growth was seen in follow-up imaging.

### 6.5 DNA sequencing

For each tumor sample with sufficient DNA yield, at least 550 µg DNA was used for DNA-sequencing. Genomic libraries were sequenced with 50-bp single-read shallow whole-genome sequencing (WGS) performed on a NextSeq500 (Illumina, San Diego, California). Sequence reads were aligned against the reference genome (GRCh37/hg19) with Burrows-Wheeler Alignment tool[53] (bwa aln; version 0.5.9) and deduplicated with Picard tools[54] (version 1.61). Reads with mapping quality *<*37 were excluded from further analysis. Libraries could be prepared for 219 out of 234 tumor samples, originating from 103 patients, a complete tumor pair was available for 93 patients. Samples with sufficient DNA after hybridization (*>*500 ng) proceeded to whole-exome sequencing (WES).

Copy-number profiles were generated with the QDNAseq package[55] (version 1.12.0), using 100-kbp bins for general copy-number analysis and 15-kbp bins to call focal gains and deletions. Segmentation was performed using the DNAcopy package[56] (version 1.50.1), cellularity and ploidy were estimated with the ACE package[57] (version 1.9.3) and copy-number calling was performed with the CGHcall package[58] (version 2.38.0). Fits were manually assessed when the estimated ploidy was *>*2n or the estimated cellularity was *>*95%. Alternative fits were visually inspected and the VAF of *IDH1* was used to estimate the cellularity as a guide to choose the most likely fit.

Copy-number heterogeneity (CNH) was determined as described by van Dijk et al.[12] based on the copy-number profiles, with the estimated cellularity and ploidy as input measures. Aneuploidy score was determined based on the segments data. The weighted median of each chromosome arm was calculated with the absolute copy number as value and the segment length (bp) as weight. Aneuploidy of a chromosome arm was called when the weighted median deviated *>*0.5 from the estimated ploidy. The total number of aneuploid arms is the per-sample aneuploidy score.

Differential copy-number analysis of chromosomal regions was performed by identification of start and end sites of all chromosomal segments, occurring through all 219 tumor samples, to create uniform breakpoints. The mean normalized log2 read count and standard error (SE = SD/sqrt(n)) for each interval was calculated per group. A Wilcoxon signed-rank test was performed to identify regions that were significantly different between initial and recurrent tumors (BH-corrected, FDR *<*0.05).

Genomic libraries were prepared according to the manufacturer’s protocols (KAPA Library Preparation, KAPA Biosystems, Wilmington, MA) WES libraries were prepared using the KAPA HyperExome Prep Kit according to the manufacturer’s protocols and sequenced on an Illumina NovaSeq 6000 machine. Libraries could be prepared for 212 tumor samples, originating from 103 patients, a complete tumor pair was available for 88 patients.

Trimming was performed with cutadapt[59], reads were aligned with bwa (version 0.7.2) and duplicates were removed with gatk::MarkDuplicates[54] (version 4.0.1.2). Calling was done with recommended settings for both Lofreq[60] (version 2.1.3.1) and Mutect2, combined with an in-house ‘panel-of-normals’, generated from 247 normal samples processed identically to the tumor samples. Cut-off for inclusion in the ‘panel-of-normals’ was presence of the mutation at a minimum variant allele fraction (VAF) of 0.0025 in at least two normal samples. Only variants called by both Mutect2 and Lofreq were included in subsequent analysis. In addition, synonymous mutations, mutations annotated as germline by Mutect2, mutations present in the Hartwig Medical Foundation Panel of Normals, and mutations with GermQ *<*2 were filtered. Tumor mutational burden was calculated by dividing the per-sample total mutation count after filtering by the size of the sequencing target region (43Mb). Cut-off for hypermutation was set at the elbow of the ordered log(TMB) curve of all WES samples. The cohort-specific hypermutation cut-off was determined to be *≥*6 muts/Mb (**Supplementary Fig. 1d**).

### 6.6 DNA-methylation profiling

For each tumor sample with sufficient DNA yield, 80-250 µg genomic DNA was bisulfite converted using the EZ DNA-methylation Kit (Zymo Research) and processed on Infinium MethylationEPIC BeadChip arrays (Illumina) according to the manufacturer’s instructions. Array data (IDAT files) were processed using the Bioconductor minfi package[61] (version 1.36.0) to obtain the raw signal intensities. Background normalization was performed using the Noob preprocessing method for Infinium methylation micro arrays, which accounts for technical variation in background fluorescence signal. Failed CpG positions were detected by comparing the total signal (M + UM) to the background signal. Tumor samples of which more than 5% of all positions were uninformative, did not pass quality control. Out of the 312 arrays that were performed, data of 235 tumor samples originating from 103 patients, were included for further analysis.

Copy number data were generated from the DNA-methylation array data using the conumee package62. Copy number data generated by sWGS and the methylation arrays were combined to call focal gains and deletions. Focal copy-number alterations were called at ±1 normalized log2 read counts on sWGS data and ±0.2 log2 intensity differences on DNA-methylation array data. Copy-number calls of *CDK4*, *CDK6*, PDGFRA, *CDKN2A/B*, MET, MYB, MYC and RB1 were visually inspected for CNA calling. Homozygous deletion was called when a clear double loss of the locus was visible, amplification was called when the gain was clearly focal and gained more than twice. Total CNV-load was calculated by summing the length of all segments with ±0.1 log2 intensity differences.

Failed positions, positions mapping to the X and Y chromosomes, and positions with known poor quality were excluded from all DNA-methylation analyses, leaving 656517 probes for DNA-methylation analysis throughout the genome[63]. The M-value of each probe is calculated as the log ratio of the intensities of the methylated versus the unmethylated probe (log(M/UM)). When more molecules are methylated than unmethylated, the M-value of the probe will be positive, while a negative value means that more molecules were unmethylated. For all analyses the M-values were used, since they are more statistically valid for differential methylation analysis than *β*-values. Quality control and preprocessing, of the MethylationEPIC BeadChip array data from 432 adult IDH-mutant astrocytoma (grade III) patients from the CATNON randomized phase 3 trial that was used as a validation set, was performed similarly.

Absolute tumor purity was predicted from DNA-methylation data using a pre-trained random forest regression model using the RFpurify package. Differentially methylated positions (DMPs) were identified with various linear regression models, using the Bioconductor limma package[64] (version 3.28.14). In the model to identify DMPs between initial and recurrent tumors, we accounted for the individual patient effect. We fitted the model, created a contrast matrix to specifically compare the difference between initial and recurrent tumors, computed the estimated coefficients and standard errors for this contrast, and used an empirical Bayes method to rank the probes in order of evidence for differential expression. Differentially methylated regions (DMRs) between initial and recurrent tumors were identified using the Bioconductor DMRcate package[65] (version 2.12.0), in combination with the same model and contrast matrix used for the DMP analysis. These regions are based on the presence of *≥*2 differentially methylated positions (DMPs) per 1,000 nucleotides. To identify DMPs between treated and untreated tumors, we made use of a mixed effects model and accounted for the individual patient effect and the possible bias of the initial tumors, as described by Law et al. (section 7.6)[66].

### 6.7 RNA sequencing

For each tumor sample with sufficient RNA yield, at least 200 µg RNA was used for RNA-sequencing. Quality and RNA concentration were determined using the Fragment Analyzer, and proceeded with samples with a DV200 *>*15%. The NEBNext Ultra II Directional RNA Library Prep Kit for Illumina was used to process the samples. The sample preparation was performed according to the protocol “NEBNext Ultra II Directional RNA Library Prep Kit for Illumina” (NEB #E7760S/L). Briefly, rRNA was depleted from total RNA using the rRNA depletion kit (NEB #E6310). After fragmentation of the rRNA reduced RNA, a cDNA synthesis was performed. This was used for ligation with the sequencing adapters and PCR amplification of the resulting product. Clustering and DNA sequencing using the NovaSeq6000 was performed according to manufacturer’s protocols, with a concentration of 1.1 nM of DNA. These experiments were performed at the GenomeScan B.V. (Plesmanlaan 1d, 2333 BZ, Leiden, The Netherlands). Paired-end raw sequencing reads were obtained from GenomeScan as compressed raw FASTQ files. Reads were trimmed with fastp and discarded based on quality, aligned with STAR 2.7.3a to GRCh38 and Gencode 34, and duplicate reads were marked with sambamba[67–69]. Read counts were obtained with featureCounts, and serve as the primary input for all bulk RNA sequencing analyses in this manuscript[70].

Data of tumor samples with less than 750.000 reads were considered of insufficient resolution for further analysis and therefore excluded. Low-count genes with an average expression of *<*3 reads per sample were removed from the data set. Median-of-ratios normalization was performed using the DESeq2 package[71] (version 1.36). For visualization and correlation analysis, read counts were transformed into pseudo-normal expression values using the Variance Stabilizing Transformation (VST) from the DESeq2 package. Out of the 217 tumor samples with available RNA-sequencing data, 169 passed quality control. Of 59 patients a complete tumor pair was available, the data originating from the 32 patients of which either the data from the initial or the recurrent tumor passed quality control, was used in further analyses to maximize statistical power.

Differential expression analysis between initial and recurrent tumors, was performed using DESeq2. To correct for the individual patients, a unique patient identifier was used, and patients without a matching pair, were combined in a separate group. To test for the influence of external factors, the DESeq2 test was corrected for various factors, and the contribution of each factor per gene was calculated. Recursive correlation clustering was performed on the identified differentially expressed genes (DEGs) using the recursiveCorPlot package[11]. The value of the first principal component (PC1), of each expression cluster found with the recursive correlation clustering, was used as a per-sample expression cluster value. Samples were divided into high and low cluster expression classes based on the per-cluster PC1 value (‘high’: PC1 *>*0; ‘low’: PC1 *<*0).

### 6.8 Single-nucleus/cell RNA-sequencing

Cryopreserved matched initial and recurrent tumor samples from three patients with CNS WHO grade 4 recurrences were selected from the GLASS-NL cohort for single-nucleus RNA-sequencing (snRNA-seq). Nuclei were isolated as described previously[11]. In brief, 0.1g of cryopreserved tumor tissue was suspended in lysis buffer (EZ lysis buffer (Sigma, Darmstadt, Germany, N3408) with 2 U/ml RNAse inhibitor (RNAseOUT; Invitrogen, Waltham, MA, USA, #10777019)) in an all glass tissue grinder (Kimble, #885300-0002) and homogenized using the two pestles to release the nuclei into the suspension. Nuclei were then filtered through a 70µm strainer mesh (Corning, #431751) and subsequently incubated in lysis buffer and washed with wash and resuspension buffer (WRB) (1x PBS with 1% BSA, 2U/ml RNAse inhibitor (Invitrogen, #10777019) and Hoechst (Sigma, #H3570)), and filtered through a 40µm strainer mesh (Corning, #431752), to remove remaining cell debris. Nuclei were immediately sorted with FACS for quality control and counting and intact nuclei were resuspended to a concentration of 700-1200 nuclei/µL and processed on the 10X single cell RNA sequencing platform following manufacturer protocols (CG000204 Rev D, 10X Genomics, Inc).

snRNA-seq data were preprocessed using cellranger mkfastq and cellranger count software from 10X Genomics (Pleasanton California, US). For each sample, a lower count limit per nucleus was determined and nuclei below this threshold or nuclei with a mitochondrial read fraction of *>*0.1 were excluded for further analysis. Duplicates were removed using the scDblFinder package[72]. All samples were normalized using the SCTransform V2 function from the sctransform package[73]. Data of all samples was integrated using the reciprocal PCA from the Seurat V4 package[74]. Cell types were annotated by using predefined marker gene lists. For healthy cell types, the top 500 genes were selected from a reference gene list75 and for the tumor cell subclusters, marker genes were used from Ventreicher et al.[76].

The enrichment score for expression of marker gene sets per nucleus was calculated using gene function described previously[29]. In short, for a test gene set ‘G’ a reference gene set ‘R’ is constructed. This is done by first binning all genes from the expression dataset into 30 bins according to expression level. For every gene in G, 100 random genes from its corresponding bin is added to R, resulting in R being 100 times larger than G. The difference in average expression for G and R is consequently calculated for each nucleus to provide an enrichment score for each marker gene set. Tumor cells of each snRNA sample were grouped into positive and negative fractions based on the enrichment scores (**Supplementary Fig. 2c**).

The pooled single-cell RNA-sequencing dataset comprises of the IDH-mutant sample [PJ016/GSE103224] from Yuan et al.[26], the IDH-mutant sample [SF11136/GSE138794] from Diaz et al.[27], and the samples annotated as “IDHmut noncodel” from Johnson et al.[28]. All samples were individually processed using the default Seurat V4 pipeline. For the scRNA samples from Yuan and Diaz, cell types were annotated in conjunction with markers from McKenzie et al.[75] and Venteicher et al.[76], the data from Johnson et al. already provided cell type annotations. Then for each of the four datasets individually (van Hijfte, Yuan, Diaz & Johnson), the average expression of the 604 DEGs per single cell/nucleus cluster was plotted using Seurats DotPlot function (col.min: -5.6, col.max: 5.6). From these DotPlots the most relevant (i.e. highest expressed) clusters were exported and included as annotation bars to the co-expression clustering.

### 6.9 Proteomics

Two 10µm FFPE tissue sections from available tumor samples were used for proteome analysis. Sample processing, data acquisition and analysis were performed at the Functional Genomics Center Zurich (FGCZ) with support from Dr. Sibylle Pfammatter, as described by Bueler et al.[77]. In short, samples were run on a Bruker timsOF attached to an EvoSep system using diaPASEF[78]. A fractionated reference library was prepared from a sample pool and protein quantification, resulting in per-patient signal intensities for all identified proteins, was performed with Spectronaut software. Proteins quantified in less than 66% of all samples were excluded, samples with *>*30% of missing values did not pass quality control and were excluded for further analysis. Data was normalized by median absolute deviation (MAD) normalization and values are imputed using a mixed imputation approach[79]. 55 out of 76 tumor samples, of which libraries could be prepared, also passed quality control. Of 18 patients a complete tumor pair was available, the data originating from the 18 patients of which either the data from the initial or the recurrent tumor passed quality control, was used in further analyses to maximize statistical power.

Differentially expressed proteins (DEPs) were identified with a linear regression model, using the Bioconductor limma package[64]. We fitted the model, created a contrast matrix to specifically compare the difference between CNS WHO grade 2&3 and CNS WHO grade 4 tumors, computed the estimated coefficients and standard errors for this contrast, and used an empirical Bayes method to rank the proteins in order of evidence for differential expression.

### 6.10 Survival analyses

All patients were followed until death or censored at date of last follow-up. Overall survival (OS) was measured from the date of initial surgery until date of death or censor ship. Progression-free survival (PFS) was measured from date of surgery until date of next progression, defined by clinical decline and/or radiological progression. Time-to-recurrence (TTR) was measured from date of initial surgery to date of recurrent surgery. Post-recurrent survival (PRS) was measured from date of recurrent surgery until date of death or censor ship. Univariable and multivariable survival analyses were performed using Cox proportional-hazards models to calculate hazard ratios (HR), and Kaplan Meier curves were generated using the survival and survminer packages[80,81].

### 6.11 Statistical methods

All statistical analyses were performed using R (version 3.5.3) and RStudio (version 1.2.1335). Plots were generated using base R and tidyverse.

## 8 Supplementary Figures

**Supplementary Fig. 1:**
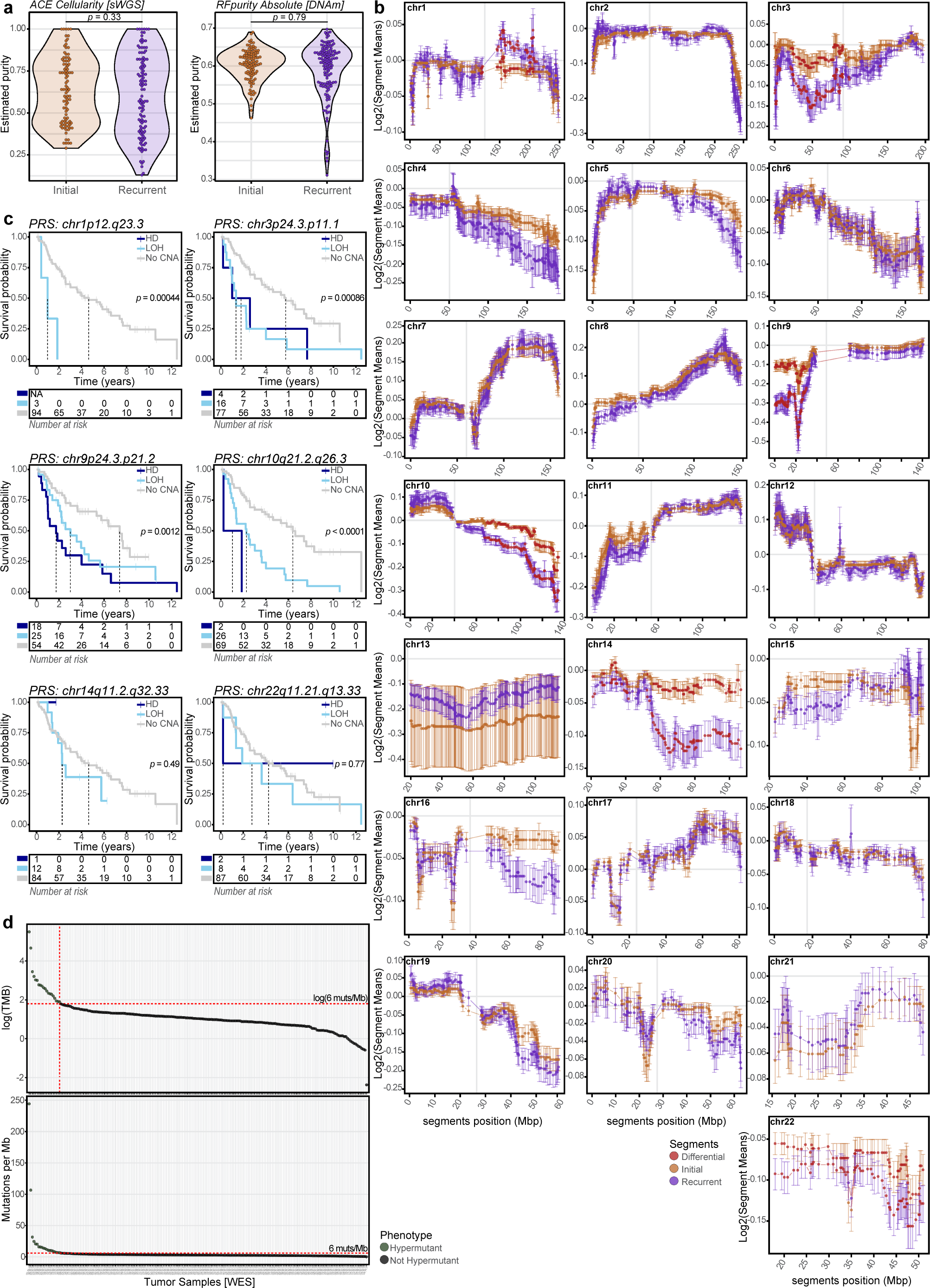
Longitudinal genomic changes. **a**, Difference in tumor purity between initial and recurrent tumor samples. Difference between initial and recurrent tumor samples is calculated using a Wilcoxon rank sum test for the ACE estimated cellularity based on sWGS data (left, *p*= 0.33) and the RFpurify Absolute estimated purity based on DNA-methylation array data (right, *p*= 0.79). **b**, Differential CNA analysis between matched initial and recurrent tumors. Each panel represents a chromosome, with on the x-axis the chromosomal position of each segment breakpoint and on the y-axis the log2(read counts), centromere is allocated with a vertical grey line. Segments mean with error bars (SE = SD/sqrt(n)) are separately visualized for initial (orange) and recurrent tumor samples (purple). Significant differential segments between initial and recurrent tumors (in red) were identified using a paired Wilcoxon signed-rank test (BH-corrected, FDR *<*0.05). Chromosomal regions chr1p12-q23.3, chr3p24.3-p11.1, chr9p24.3-p21.2, chr10q21.3-q26.3, chr14q11.2-q32.33, and chr22q11.21-q13.33 had significantly lower log2 read counts in recurrent tumors compared to initial tumors. **c**, Kaplan-Meier curves for PRS when stratifying patients on the CNA status (HD *<*-0.4 and LOH *<*-0.2 mean log2(read counts)) of differential regions (identified in Extended Data Fig. 1b) of the recurrent tumor. Compared to cases without an alteration, loss of chr3p24.3.p11.1 (LOH (n = 16): *p*= 7.74*10*^−^*^4^, HR = 2.900[1.559-5.395]; HD (n = 4): *p*= 0.075, HR = 2.564[0.9101-7.223]), chr9p24.3-p21.2 (LOH (n = 25): *p*= 0.024, HR = 2.035[1.098-3.770]; HD (n = 18): *p*= 6.57*10*^−^*^4^, HR = 3.106[1.618-5.963]) and chr10q21.3-q26.3 (LOH (n = 26): *p*= 8.62*10*^−^*^5^, HR = 2.968[1.724-5.109]; HD (n = 2): *p*= 0.004, HR = 8.888[2.048-38.582]) were negatively associated with PRS. **d**, Hypermutant cut-off value for the GLASS-NL cohort. Upper panel shows the log scale of the TMB (muts/Mb) of each tumor sample, ordered based on the log(TMB). The elbow of the curve is located at 6 mutations per Mb and marked with a horizontal red dashed line. Samples located on the left side of the vertical red dashed line are considered to be hypermutated. Lower panel depicts the TMB per sample with the cut-off of 6 muts/Mb marked with a red dashed line. Hypermutant samples are colored in green. sWGS = shallow whole-genome sequencing, CNA = copy-number alteration, PRS = post-recurrent survival, HD = homozygous deletion, LOH = loss of heterozygosity, TMB = tumor mutational burden.

**Supplementary Fig. 2:**
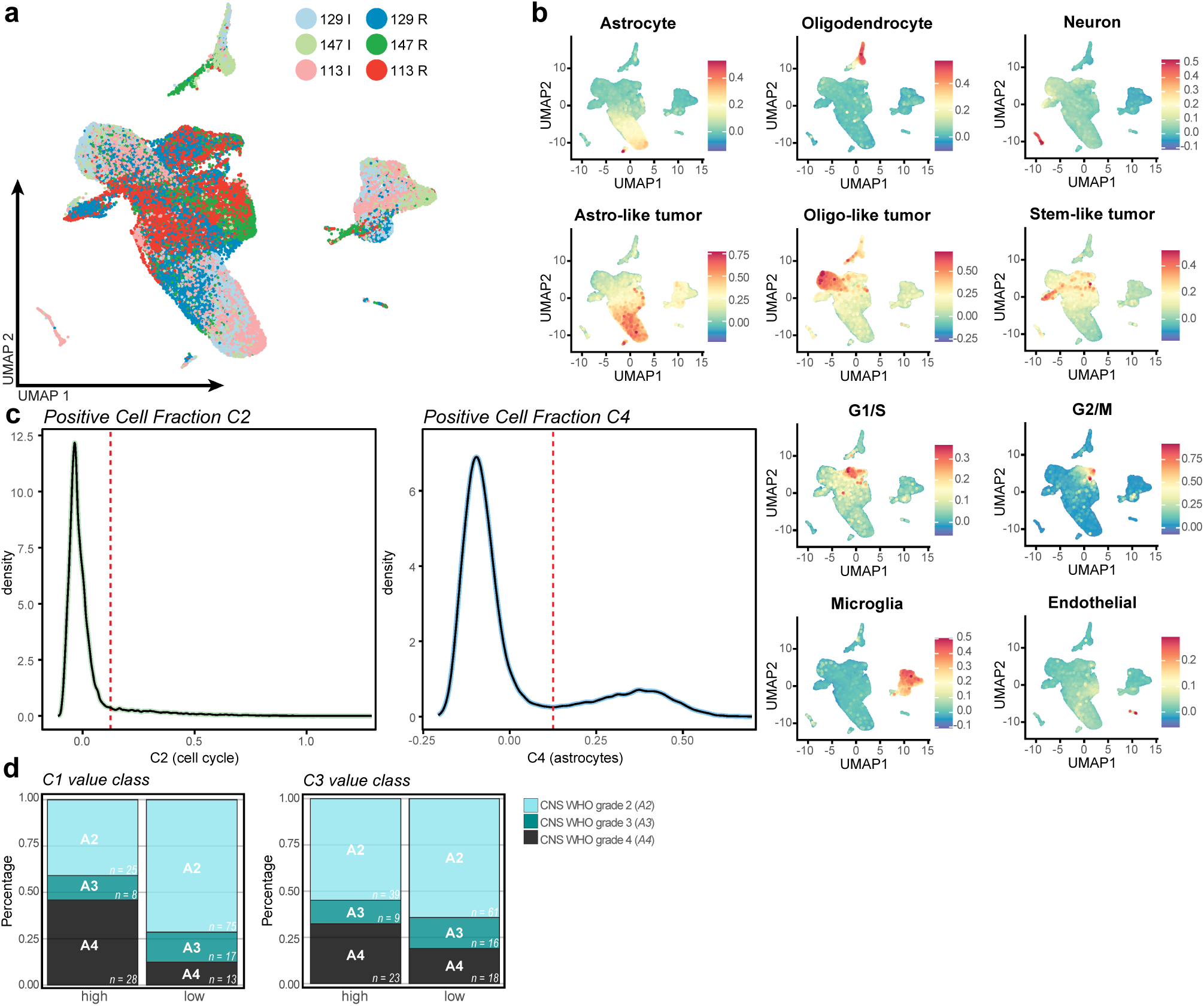
snRNA validation of gene expression clusters. **a**, The integrated snRNA dataset of all snRNA samples (n = 6) from three IDH-mutant astrocytoma patients from the GLASS-NL cohort. **b**, UMAP projection of single nuclei of integrated dataset, colored by the enrichment score of the cell type clusters. **c**, Density plots of expression of gene expression cluster C2 (left) and C4 (right) by single nuclei. Dashed line indicates cutoff for cells to be marked as ‘positive’ cluster expression. **d**, Distributions of CNS WHO grades within the expression cluster value classes (‘high’: PC1 *>*0; ‘low’: PC1 *<*0) of C1 and C3 (graphs of C2 and C4 are found in **Fig. 5d**).

